# SnakeHichipTF reveals transcription factor logic underlying enhancer-promoter wiring in the human brain

**DOI:** 10.64898/2026.03.30.715320

**Authors:** Jiang Tan, Yuqing Wu, Richard Head, Yidan Sun

**Affiliations:** Department of Genetics, Washington University School of Medicine, St. Louis, Missouri, USA; Center for Translational Bioinformatics, Washington University School of Medicine, St. Louis, Missouri, USA; McDonnell Genome Institute, Washington University School of Medicine, St. Louis, Missouri, USA; Institute for Informatics, Data Science and Biostatistics (I2DB), Washington University School of Medicine, Saint Louis, Missouri, USA

**Author notes:** These authors contribute equally (Jiang Tan, Yuqing Wu).

**Keywords:** HiChIP, enhancer-promoter interactions, 3D genome organization, transcription factor footprinting, Middle Frontal Gyrus, Substantia Nigra, Human Accelerated Regions, neuropsychiatric traits, chromatin architecture

## Abstract

Enhancer-promoter interactions are a central feature of gene regulation, yet the regulatory logic that governs their selective formation in complex tissues remains poorly understood. To address this gap, we developed SnakeHichipTF, a reproducible and scalable framework that integrates multi-engine HiChIP analysis with AI-based footprinting to decode the transcription factor (TF) logic underlying enhancer-promoter wiring. Applying SnakeHichipTF to HiChIP datasets from the human Middle Frontal Gyrus (MFG) and Substantia Nigra (SN), we identified distinct region-biased enhancer interactions associated with differential gene expression. MFG-biased interactions were enriched for cognitive and psychiatric associated GWAS traits, whereas SN-biased interactions preferentially intersected lipid and metabolic trait architectures. Integration of TF footprinting revealed that these region-biased interaction networks are governed by distinct TF programs: MFG-biased interactions preferentially recruited TFs linked to neuronal signaling and transcriptional activation, whereas SN-biased interactions were associated with metabolic and stress-responsive regulators. Interestingly, MFG-biased regulatory interactions were significantly enriched for Human Accelerated Regions (HARs), and HAR-associated genes showed elevated expression in humans relative to non-human primates, indicating that cortical enhancer wiring is embedded within evolutionarily modified regulatory elements. Together, by linking 3D chromatin architecture, TF logic, genetic risk, and evolutionary regulatory elements, SnakeHichipTF provides a general framework for dissecting the mechanistic basis of spatial gene regulation.

## Introduction

Long-range enhancer-promoter interactions are a fundamental organizing principle of gene regulation, enabling distal regulatory elements to activate target genes in a cell-type–specific manner [1–5]. Advances in chromatin conformation assays [6,7] have revealed that enhancers and promoters frequently engage in three-dimensional (3D) contacts that correlate with transcriptional activity, developmental programs, and disease-associated regulatory variation [1–5]. However, despite the widespread presence of active enhancers and promoters across the genome, only a subset of potential enhancer-promoter pairs form stable 3D interactions. What determines the selective formation of enhancer-promoter loops in complex genomes remains a fundamental and unresolved question in gene regulation.

One emerging view is that enhancer-promoter loop formation is not merely a passive consequence of genome folding, but an actively regulated process orchestrated by transcription factors (TFs) [8–11]. In a broad sense, TFs encompass a spectrum of DNA-binding proteins, ranging from sequence-specific transcription factors to architectural regulators such as CTCF and YY1 [12]. By recognizing DNA motifs and recruiting cofactors [12], including cohesin, BRD4, and the Mediator complex, these proteins help establish and stabilize enhancer-promoter interactions through mechanisms such as loop extrusion [13], condensate formation[8], and coordinated chromatin remodeling. Distinct cell types and tissues express unique repertoires of TFs [14], suggesting that context-specific TF combinations may function as regulatory assembly codes that determine which enhancers engage their target promoters in three-dimensional space. However, it remains unclear which cooperative TF programs selectively drive enhancer-promoter loop formation in specific cellular contexts and how such programs translate into robust gene expression outputs.

Progress has been limited by the absence of systematic frameworks to interrogate the TF regulatory logic encoded within enhancer-promoter interaction maps. While HiChIP and PLAC-seq [6,7] enable genome-wide mapping of enhancer interactions, current analyses largely treat detected loops as structural endpoints rather than components of coordinated regulatory programs. Conversely, chromatin accessibility assays such as ATAC-seq, combined with footprinting-based inference [15,16], can profile TF binding activity at scale but lack direct information about distal enhancer-promoter connectivity. As a result, TF-centric and 3D interaction–centric analyses remain largely disconnected, constraining mechanistic interpretation of enhancer-promoter loops.

To bridge this gap, we developed SnakeHichipTF, an integrated computational framework designed to interrogate the regulatory logic underlying enhancer-promoter loop formation. SnakeHichipTF unifies multiple widely used HiChIP interaction-calling strategies within a modular and reproducible workflow, enabling standardized processing of chromatin interaction data while preserving method-specific variation. Crucially, rather than treating interaction calls as static endpoints, the framework couples multi-engine upstream interaction detection with a downstream AI-based ATAC-seq footprinting module for systematic TF inference. By directly integrating TF binding activity with enhancer-promoter interaction maps, SnakeHichipTF provides a principled strategy to decode the TF programs that govern selective enhancer-promoter loop formation.

By applying SnakeHichipTF to published H3K27ac HiChIP datasets from distinct human brain regions, we identified region-specific enhancer-promoter interactions that were selectively enriched within distinct brain compartments. Integration with an AI-based ATAC-seq footprinting module further revealed that these region-specific interaction landscapes were associated with distinct TF programs, many of which are known to regulate corresponding neurobiological functions. These findings support a model in which the selective formation of enhancer-promoter loops reflects combinatorial TF activity rather than stochastic chromatin contacts. Together, our results establish SnakeHichipTF as an end-to-end framework that directly connects transcription factor activity to 3D chromatin architecture and deciphers the regulatory logic governing enhancer-promoter interactions.

## Results

### SnakeHichipTF provides a unified framework for HiChIP data analysis and transcription factor inference

SnakeHichipTF is organized as a modular framework that connects genome preprocessing, HiChIP interaction detection, and TF inference into a unified workflow (Fig. 1A). The pipeline is organized into three coordinated modules designed to ensure reproducibility, analytical flexibility, and mechanistic interpretability.

**Figure 1.**
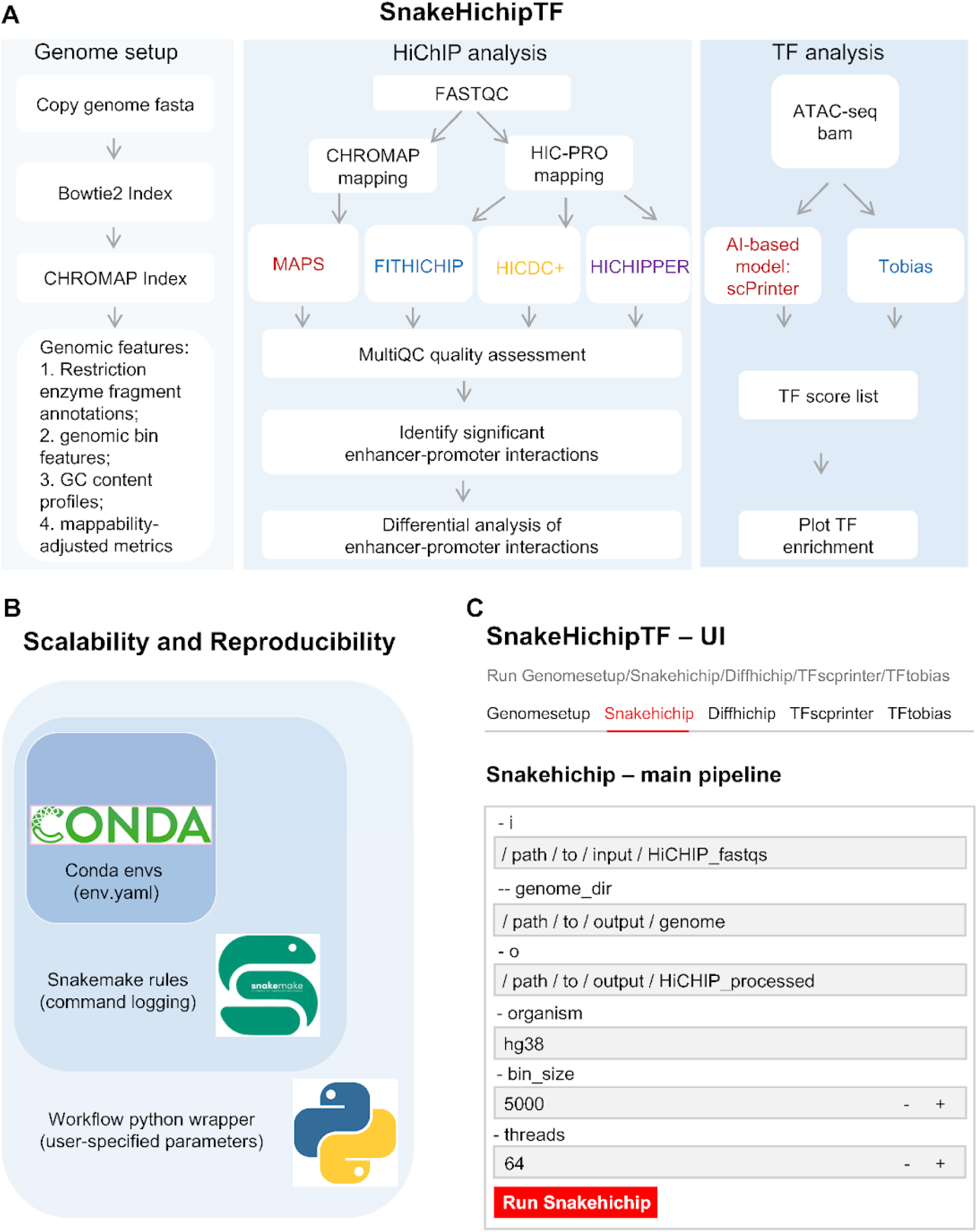
SnakeHichipTF provides a unified framework for HiChIP data analysis and transcription factor inference. Schematic overview of the SnakeHichipTF pipeline. The workflow is organized into three top-level modules: Genome setup, HiChIP analysis, and transcription factor (TF) analysis. **(A)** The genome setup module prepares standardized reference files, including CHROMAP and Bowtie2 indices, alongside customized genomic features (restriction enzyme fragment annotations, genomic bin features, GC content profiles, and mappability-adjusted metrics). The HiChIP analysis module processes raw data through mapping (CHROMAP and HIC-PRO) and quality assessment (FASTQC and MultiQC). Significant enhancer-promoter interactions are identified using a multi-engine core comprising FITHICHIP, MAPS, HICDC+, and HICHIPPER, enabling robust differential analysis. The TF analysis module integrates ATAC-seq data using Tobias and the AI-based model scPrinter to generate TF score lists and enrichment plots. **(B)** Scalability and reproducibility are ensured through three integrated layers: Conda environments isolate and fix software versions across systems; Snakemake rules record every execution step and manage command logging and a Python workflow wrapper standardizes user-specified parameters. **(C)** A Streamlit-based graphical user interface (Local UI) provides a user-friendly dashboard for pipeline configuration and execution, enhancing accessibility for users without extensive command-line expertise.

The workflow initiates with a comprehensive Genome Setup module, ensuring that all subsequent analyses are anchored to a consistent genomic coordinate system. This module automates the generation of essential genomic resources, including reference genome indexing, restriction fragment site identification, and chromosome size calculation (Fig. 1A, left panel). By standardizing these prerequisites, SnakeHichipTF minimizes cross-study variation and supports seamless deployment across diverse species.

At the core of the framework is the HiChIP module for identifying differential enhancer-promoter interactions. Sequencing reads undergo quality control and mapping using CHROMAP [17] or HiC-Pro [18], followed by interaction calling with four widely adopted HiChIP-specific engines: MAPS [7], FitHiChIP [19], HiC-DC+ [20], and hichipper [21] (Fig. 1A, center panel). These tools were selected because they were specifically developed to model enrichment-based chromatin interaction profiles characteristic of HiChIP and PLAC-seq datasets. By integrating multiple engines rather than relying on a single algorithm, SnakeHichipTF accommodates variation in sequencing depth, enrichment strength, and experimental design. This multi-engine strategy reflects the predominant analytical practices in the field and allows users to select the most appropriate method for their datasets while preserving a harmonized downstream representation. Significant enhancer-promoter interactions are subsequently identified and subjected to differential analysis, enabling systematic characterization of condition-specific enhancer-promoter regulatory landscapes.

The final module links enhancer-promoter interactions to underlying transcription factor activity (Fig. 1A, right panel). We integrated two complementary TF inference approaches: scPrinter [16], an AI-based model that captures context-dependent TF activity patterns, and TOBIAS [15], a bias-corrected footprinting framework grounded in established accessibility modeling. The inclusion of both methods reflects a balance between model-driven predictive inference and classical footprint-based estimation, enabling robust TF activity quantification across datasets with varying signal complexity. TF activity scores are then mapped onto corresponding enhancer-promoter interactions, allowing systematic association of differential enhancer-promoter interactions with underlying transcriptional programs. This integrative design enables mechanistic interpretation of how combinatorial TF activity contributes to the selective formation of enhancer-promoter loops.

### Modular architecture of SnakeHichipTF ensures scalable and reproducible analysis

Beyond simple tool aggregation, SnakeHichipTF reconstructs the original analytical logic of each engine within a reproducible Snakemake architecture. Early HiChIP tools were implemented as standalone bash workflows with implicit preprocessing assumptions and fragmented execution steps. We systematically decomposed these pipelines, preserved their recommended mapping and normalization strategies, and reimplemented them as modular, logged, and environment-controlled Snakemake rules. Snakemake automatically records all input files, parameters, intermediate steps, and outputs, generating a complete execution graph and audit trail. In parallel, all software dependencies and version specifications are locked within Conda environment files to ensure consistent execution across systems. Together, explicit environment management and automated provenance tracking ensure full reproducibility of analytical results (Fig. 1D).

This architecture also supports efficient scaling across diverse computational environments, from local workstations to high-performance computing clusters. The workflow parallelizes computationally intensive steps, including alignment, interaction calling, and differential analysis, across samples and engines. On a 32-core workstation, processing a 50-million-read paired-end HiChIP dataset required approximately 2–3 hours for full interaction detection and downstream integration, with near-linear acceleration when multiple libraries were processed concurrently. Memory usage remained moderate (<40 GB), demonstrating suitability for large-scale enhancer-promoter analyses across multiple conditions or tissues.

SnakeHichipTF is distributed as a standalone package via Conda and PyPI. A Python-based wrapper standardizes parameter specification and execution, enabling the entire workflow to be launched through a single command-line interface while minimizing manual configuration errors. In addition, an optional Streamlit-based graphical interface allows users to upload HiChIP data, configure analysis parameters, and adjust statistical thresholds such as P-value (Fig. 1E). Together with Snakemake’s compatibility with common job schedulers (e.g., SLURM), these features enable deployment across diverse computational infrastructures while remaining accessible to users with varying levels of computational expertise.

### Distinct enhancer-promoter wiring programs emerge across functional axes of the human brain

To investigate how enhancer-promoter wiring contributes to the functional specialization of the human brain, we focused on the Middle Frontal Gyrus (MFG), a cortical region central to higher-order cognition, and contrasted it with the Substantia Nigra (SN), an evolutionarily conserved midbrain structure primarily involved in motor control. These regions represent distinct functional and evolutionary axes of brain organization, providing a biologically meaningful contrast to interrogate the regulatory determinants of 3D chromatin architecture.

Unlike standard comparative approaches that often rely on the naive intersection of independently called loops, we leveraged the quantitative statistical framework HiCDC+ within SnakeHichipTF to perform a direct, locus-by-locus differential interaction analysis. This rigorous approach effectively controls for sequencing depth biases and allows for the precise identification of true region-biased chromatin contacts. In total, we identified 1,397 MFG-biased and 558 SN-biased interactions (Fig. 2A, B; Supplementary Table 1). The higher prevalence of MFG-biased interactions suggests a more expansive and complex enhancer-driven regulatory architecture within the prefrontal cortex compared to the midbrain.

**Figure 2.**
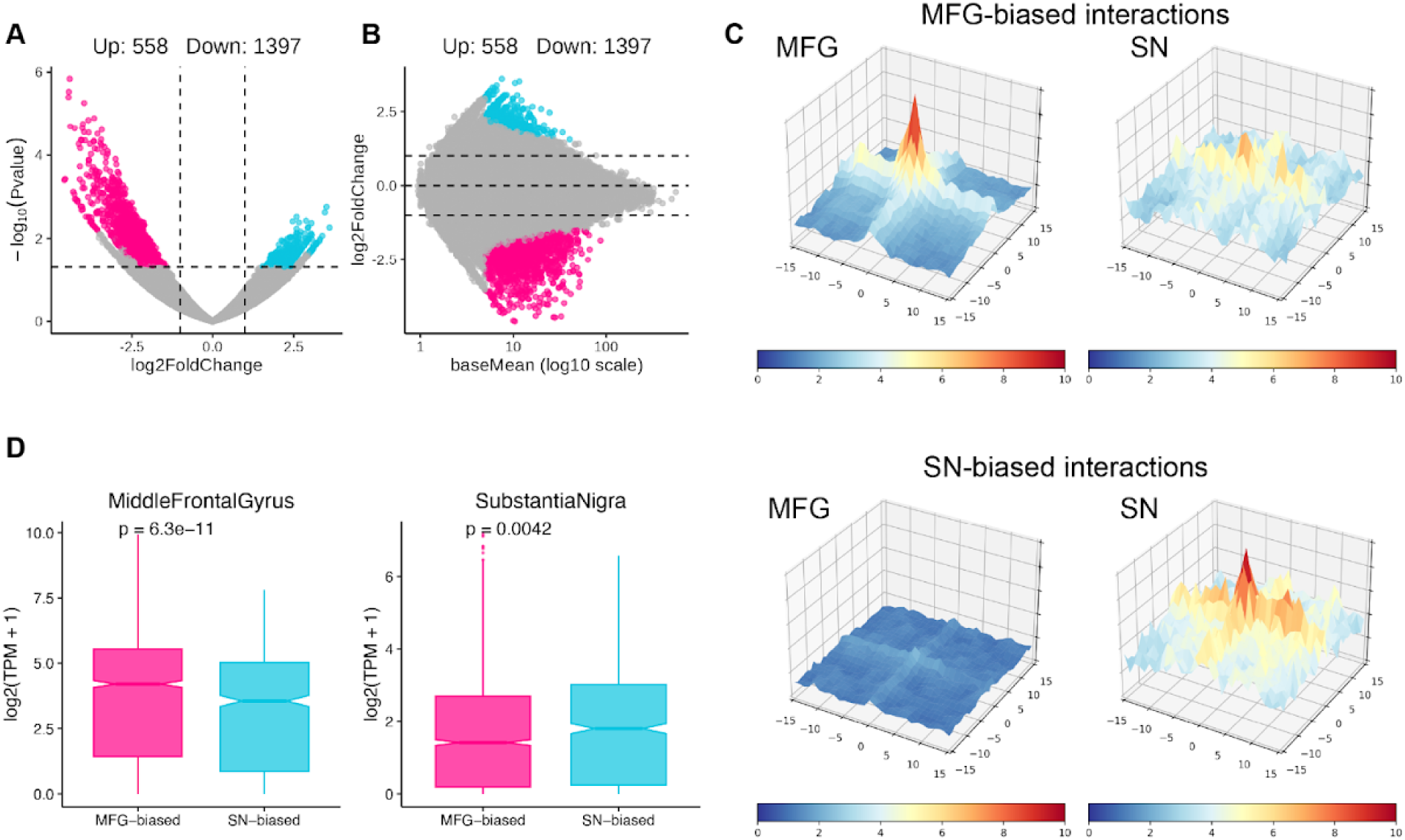
Distinct enhancer-promoter wiring programs emerge across functional axes of the human brain. **(A)** Volcano plot and **(B)** MA plot illustrating the differential chromatin interactions identified between the MFG and SN. Significant interactions are highlighted, revealing 1,397 MFG-biased (red) and 558 SN-biased (blue) enhancer interactions. **(C)** Aggregate contact maps for region-biased interaction sets. For each interaction set (MFG-biased, top; SN-biased, bottom), contacts were aggregated across all loops and plotted separately in MFG (left) and SN (right). The intensity reflects the average contact enrichment centered on loop anchors. MFG-biased interactions show stronger central enrichment in MFG than in SN, whereas SN-biased interactions show stronger central enrichment in SN than in MFG, confirming region-preferential interaction strength. **(D)** Boxplots comparing the transcript abundance (log2(TPM + 1)) of genes associated with the region-biased interaction anchors. Genes linked to MFG-biased interactions exhibit significantly higher expression in the MFG (p=6.3×10−11, left panel), whereas genes linked to SN-biased interactions display elevated expression in the SN (p=0.0042, right panel), demonstrating the functional coupling of 3D spatial wiring to region-specific transcription.

Aggregate interaction profiles further illustrated this regional asymmetry: MFG-biased interactions showed stronger and more focal interaction signals in the MFG than in the SN, whereas SN-biased interactions displayed the opposite pattern, with signal enrichment centered in the SN (Fig. 2C). These patterns support the presence of region-preferential 3D chromatin wiring rather than simple overlap differences between interaction maps.

To evaluate the functional consequences of this region-biased wiring, we integrated these spatial maps with regional transcriptomic data to examine gene expression levels associated with the interaction anchors. Genes linked to MFG-biased interactions exhibited significantly higher expression in the MFG compared to the SN (p=6.3×10−11) (Fig. 2D, left). Conversely, genes associated with SN-biased interactions showed a corresponding elevation in expression specifically within the SN (p=0.0042) (Fig. 2D, right). The concordance between 3D interaction bias and transcript abundance supports a functional coupling between enhancer-promoter wiring and region-biased gene transcription.

Collectively, these findings demonstrate that distinct enhancer-promoter wiring programs operate across the functional axes of the human brain, establishing a robust structural foundation for the subsequent dissection of the transcription factor networks that govern selective loop formation.

### Region-biased enhancer-promoter wiring intersects distinct genetic risk architectures

To further explore the biological and clinical implications of the region-biased enhancer-promoter interactions, we performed a comprehensive enrichment analysis using Genome-Wide Association Study (GWAS) loci across a diverse array of human traits and diseases[22].

An initial categorization of the enriched GWAS traits revealed a clear contrast between the two brain regions. MFG-biased interactions were predominantly enriched for “Neuro Brain” and “Proteins Level” categories, accounting for 51.5% and 33.3% of the total enriched signals (Fig. 3A). In contrast, SN-biased interactions exhibited a distinct regulatory signature, with enrichment concentrated in “Lipids Metabolites” (83.6%) (Fig. 3B).

**Figure 3.**
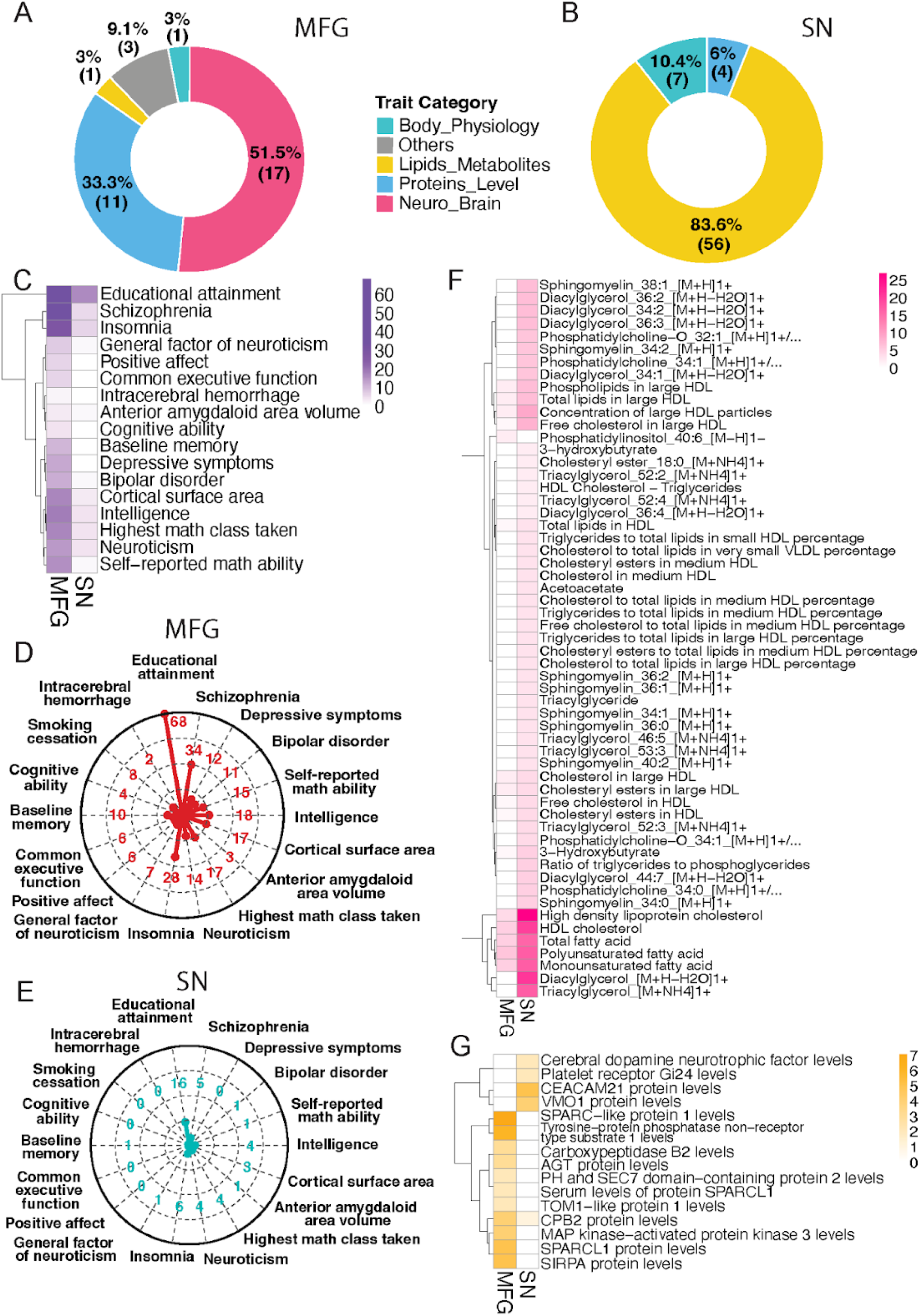
Region-biased enhancer-promoter wiring intersects distinct genetic risk architectures. **(A, B)** Distribution of enriched GWAS trait categories for **(A)** MFG-biased and **(B)** SN-biased enhancer-promoter interactions. MFG-biased interactions are predominantly enriched for “Neuro Brain” and “Proteins Level” traits, while SN-biased interactions are heavily concentrated in “Lipids Metabolites”. **(C)** Heatmap displaying the enrichment profiles of “Neuro Brain” traits. MFG-biased interactions show strong, specific associations with cognitive functions and psychiatric disorders, including educational attainment, schizophrenia, intelligence, and bipolar disorder. Color intensity represents the −log10(P-value) of the enrichment. **(D, E)** Radar plots comparing the enrichment patterns of cognitive and psychiatric traits within the **(D)** MFG and **(E)** SN interaction networks, illustrating the selective capture of neuropsychiatric genetic risk by the spatial regulatory programs of the MFG. The radial distance and corresponding numerical values indicate the GWAS number. **(F)** Heatmap detailing the enrichment of lipid and metabolite GWAS traits, highlighting significant SN-specific associations with diverse lipid species, such as sphingomyelins and diacylglycerols. Color intensity represents the −log10(P-value) of the enrichment. **(G)** Heatmap of protein-level GWAS traits distinguishing the regional profiles. MFG-biased contacts are enriched for neuronal signaling and synaptic regulation markers, whereas SN-biased contacts are enriched for circulating and midbrain-relevant proteins. Color intensity represents the −log10(P-value) of the enrichment.

Detailed examination of the “Neuro Brain” category in the MFG revealed consistent enrichment for traits associated with cognitive function and psychiatric risk (Fig. 3C). Heatmap analysis (Fig. 3C) and radar plots (Fig. 3D) showed that MFG-biased interactions were strongly associated with educational attainment, schizophrenia, intelligence, and common executive function. Traits such as bipolar disorder, depressive symptoms, and intelligence displayed prominent MFG-specific enrichment patterns (Fig. 3C). Comparative radar plots further illustrated this regional specificity: whereas the MFG interaction network captured these cognitive and psychiatric signals (Fig. 3D), the SN network showed minimal enrichment for the same traits (Fig. 3E). These results suggest that the spatial regulatory programs of the MFG preferentially intersect genetic loci underlying neuropsychiatric phenotypes.

Conversely, SN-biased interactions showed a focused enrichment for metabolic pathways. Significant associations were observed for multiple lipid species, including sphingomyelins (e.g., 38:1, 34:2) and diacylglycerols (e.g., 36:2, 34:2) (Fig. 3F). Lipid metabolism and membrane composition play essential roles in midbrain dopaminergic function and neuronal maintenance, providing a functional context for the metabolic enrichment observed in the SN. In contrast, these metabolic associations were largely absent from the MFG interaction landscape.

Protein-level GWAS traits further distinguished the regional interaction profiles (Fig. 3G). MFG-biased interactions were associated with protein markers implicated in neuronal signaling and synaptic regulation, including SPARCL1 and MAP kinase–related proteins. Meanwhile, SN-biased contacts were enriched for GWAS signals related to circulating or systemic protein levels, such as cerebral dopamine neurotrophic factor (CDNF) and CEACAM21.

Together, these results indicate functional partitioning of region-biased 3D chromatin wiring. MFG-specific regulatory interactions preferentially align with genetic architectures underlying cognitive and psychiatric traits, whereas SN-specific wiring is more closely associated with lipid and metabolic programs. This divergence provides a genetic and phenotypic context for subsequent dissection of the transcription factor programs governing selective loop formation.

### Transcription factor logic shapes region-specific enhancer-promoter wiring

To dissect the regulatory logic underlying the selective formation of enhancer-promoter loops in two brain regions, we profiled transcription factor (TF) binding within MFG-biased and SN-biased interactions using scPrinter[16], an AI-based footprinting framework applied to matched ATAC-seq data [1,5], implemented through the SnakeHichipTF pipeline.

We first quantified TF binding strength and distribution across MFG-biased and SN-biased interaction respectively (Fig. 4A, B). While some TFs exhibited widespread binding across numerous interactions, others were restricted to a small subset of region-specific contacts, indicating a heterogeneity of TF occupancy (Fig. 4C, D). For example, the architectural protein CTCF, a canonical organizer of 3D chromatin structure, exhibited high presence frequency and consistent binding scores in both the MFG and SN (Fig. 4C–F), supporting its role as a shared structural scaffold. Beyond common architectural anchors like CTCF, we identified additional TFs with high presence and binding scores that were preferentially associated with subsets of interactions (Fig. 4E, F), suggesting potential roles in region-modulated loop stabilization.

**Figure 4.**
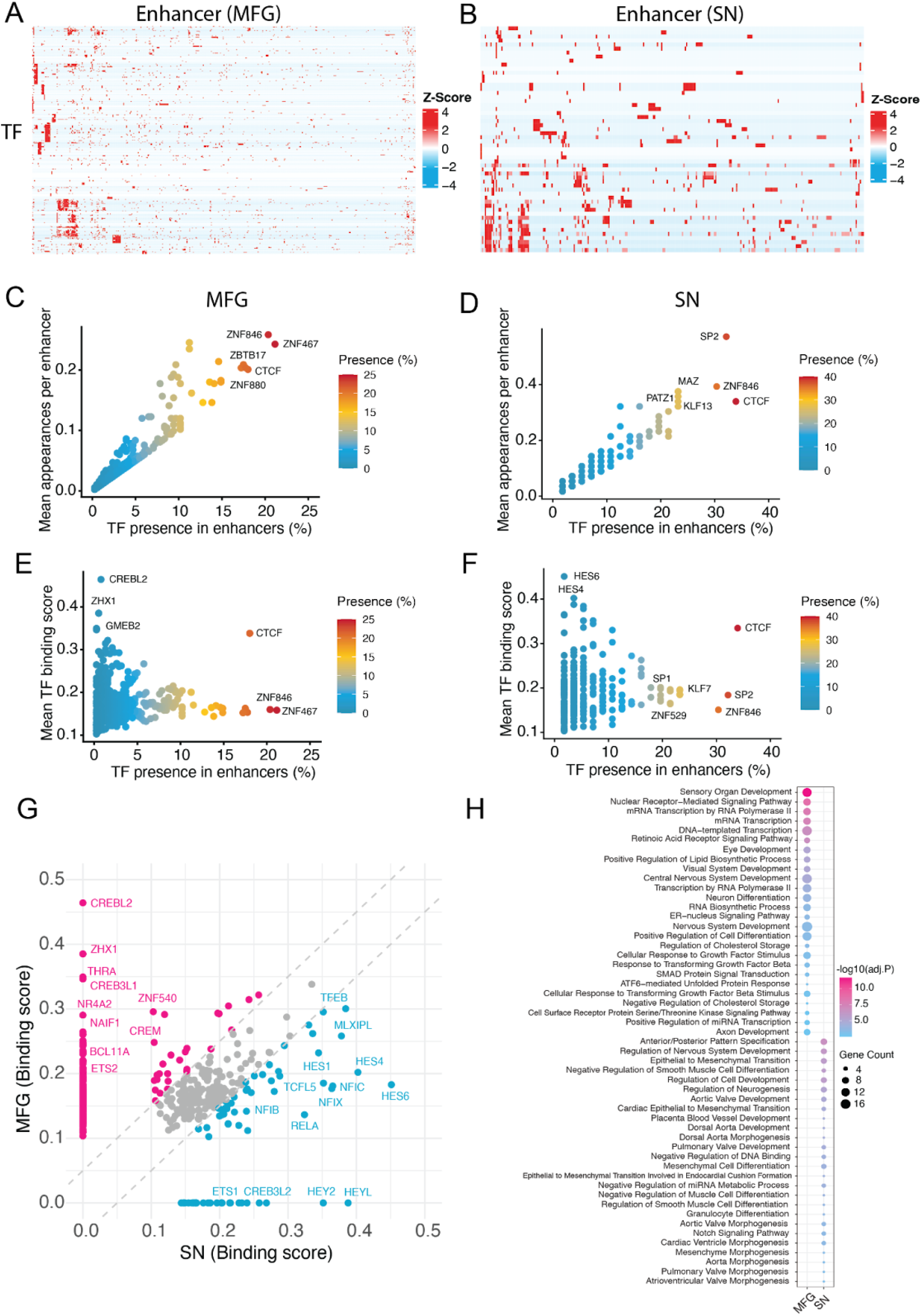
Transcription factor logic shapes region-specific enhancer-promoter wiring. **(A, B)** Heatmaps depicting the binding strength of transcription factors (TFs) across **(A)** MFG-biased and **(B)** SN-biased enhancer-promoter interactions, as inferred by scPrinter. Color represents Z-score of binding score. **(C, D)** Scatter plots illustrating the distribution of TFs across **(C)** MFG-biased and **(D)** SN-biased enhancer interactions. The y-axis represents the mean appearances per interaction, and the x-axis represents the percentage of interactions where the TF is present. **(E, F)** Scatter plots showing the mean TF binding score versus the percentage of TF presence in **(E)** MFG-biased and **(F)** SN-biased enhancer interactions. **(G)** Scatter plot correlating the binding scores of TFs between the SN (x-axis) and MFG (y-axis). TFs demonstrating significant regional bias are highlighted, including MFG-enriched factors (red) and SN-enriched metabolic regulators (blue). **(H)** Dotplot showing the top enriched Gene Ontology (GO) Biological Process terms for the region-biased TFs in the MFG and SN. The color scale represents the −log10(Adjusted P-value) and the dot size represents gene count.

To identify the TFs that specifically define the regulatory logic of each region, we compared each TF binding score between the MFG and SN. This analysis revealed distinct sets of region-biased TFs (Fig. 4G). MFG-biased interactions exhibited significantly higher binding scores for TFs such as CREBL2, ZHX1, CREB3L1, and NR4A2 (Fig. 4G). These factors have been implicated in neuronal signaling and cortical development [23–27]. Conversely, SN-biased interactions were enriched for metabolic and physiological regulators, including TFEB, MLXIPL, and members of the HES family (e.g., HES1, HES4) [28–33] (Fig. 4G).

To assess the biological relevance of these region-biased TFs, we performed Gene Ontology (GO) enrichment analysis. TFs biased toward MFG were significantly enriched for processes related to transcriptional regulation and neural differentiation, including *transcription by RNA polymerase II*, *RNA biosynthetic process*, *neuron differentiation*, *central nervous system development*, *sensory organ development*, and *axon development* (Fig. 4H). These enrichments are consistent with our earlier observations of neuron-associated enhancer interactions in the MFG and suggest that the 3D interactome in this region preferentially engages TF programs linked to transcriptional activation and neural circuit-related functions. In contrast, TFs biased toward SN were enriched for pathways associated with developmental patterning and cellular differentiation, including *regulation of neurogenesis*, *regulation of nervous system development*, *anterior/posterior pattern specification*, and *epithelial to mesenchymal transition* (Fig. 4H). Together, these results indicate that MFG- and SN-biased enhancer interactions are associated with distinct transcription factor programs, with the MFG favoring transcription-centered neural regulatory functions and the SN reflecting a more developmental and lineage-patterning-oriented regulatory landscape.

Collectively, these results suggest that region-specific enhancer-promoter wiring may be coordinated by distinct TF programs. It potentially supports a model in which transcription factor regulatory logic shapes 3D chromatin architecture in a region-dependent manner, thereby coupling enhancer-promoter interactions to distinct gene expression programs and functional specialization across the human brain.

### Human accelerated regions are preferentially embedded in MFG-biased regulatory wiring

Given the central role of the MFG in higher-order cognition and its high vulnerability to neuropsychiatric disorders, we reasoned that its underlying regulatory architecture might have been a primary target of recent human evolution. To test this, we analyzed Human Accelerated Regions (HARs), genomic elements that are highly conserved across vertebrates but have acquired a disproportionate number of human-specific mutations, often serving as the regulatory basis for human-unique traits. Given that HARs are thought to contribute to human-specific neurodevelopmental and cognitive traits, we asked whether MFG- and SN-biased interactions differentially intersect these evolutionarily accelerated elements.

We first quantified the number of HARs [34] located within MFG- and SN-biased interaction anchors. We found that MFG-biased interactions were significantly enriched for HARs (p=0.0017; Fig. 5A). In contrast, no significant enrichment of HARs was observed within the SN-biased interactions (p=0.0902; Fig. 5A). Representative genomic loci are shown to illustrate this enrichment (Fig. 5B,C). This finding suggests that the 3D regulatory landscape of the prefrontal cortex has undergone more extensive structural rewiring during human evolution compared to more conserved midbrain structures like the SN.

**Figure 5.**
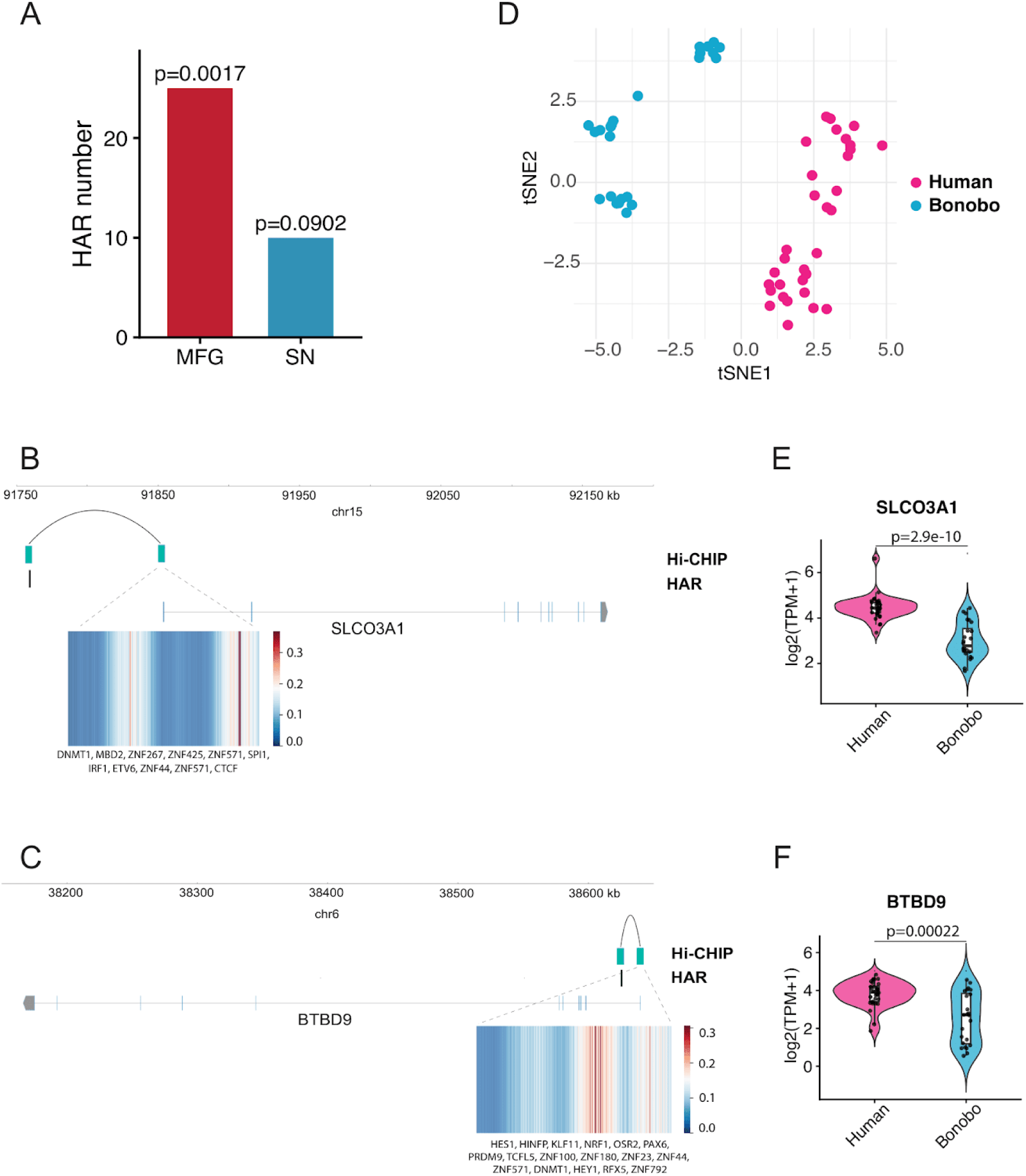
Human accelerated regions are preferentially embedded in MFG-biased regulatory wiring. **(A)** Bar chart quantifying the number of HARs located within region-biased interaction anchors. Fisher test is performed to test the significance of the enrichment. MFG-biased interactions show significant enrichment for HARs (p=0.0017), whereas SN-biased interactions do not (p=0.0902). **(B,C)** Representative genomic snapshots illustrating the integration of HARs into MFG-biased interactions for the target genes **(B)** *SLCO3A1* and **(C)** *BTBD9*. The tracks display HiChIP loops connecting HAR-overlapping distal enhancers to gene promoters. Below each track, heatmaps depict the local predicted binding affinities of specific transcription factors. **(D)** A t-SNE projection demonstrating the distinct expression divergence of HAR-regulated genes between Human (red) and Bonobo (blue) samples. **(E,F)** Violin plots comparing the expression levels (log2(TPM+1)) of HAR-regulated genes **(F)** *SLCO3A1* and **(G)** *BTBD9* between humans and bonobos. Both genes exhibit significantly higher expression in humans (**** p<0.0001, *** p<0.001), supporting the functional impact of human-specific 3D spatial rewiring.

To determine the mechanism by which these HARs are regulated, we overlapped the HAR coordinates with our previously identified TF binding profiles. Interestingly, many MFG-biased TFs showed an enrichment of binding sites within HARs (Fig. 5B,C). This suggests that the evolutionary mutations within HARs may have created novel binding platforms for “cognitive TFs,” thereby enabling the specific activation of these loops in the human MFG.

Finally, to assess whether this evolutionary rewiring resulted in functional divergence, we compared expressions of genes regulated by HAR-associated interactions between Human and another primate species Bonobo. Principal Component Analysis (PCA) demonstrated a clear separation of the human regulatory profile from bonobos (Fig. 5D). Intriguingly, certain genes, such as *SLCO3A1* and *BTBD9*, exhibited significantly higher expression levels in the human MFG compared to bonobos (Fig. 5E,F). This indicates that the acceleration of 3D genomic wiring has led to the emergence of human-specific transcriptional programs that likely underpin advanced cognitive functions.

Together, these results suggest that MFG-biased enhancer-promoter interactions not only align with cognitive and psychiatric genetic architectures but are also preferentially embedded within evolutionarily accelerated regulatory elements. The convergence of TF logic, 3D chromatin architecture, and HAR-associated expression divergence provides a potential mechanistic link between genomic acceleration and the emergence of advanced human cognitive traits.

## Discussion

In this study, we presented SnakeHichipTF, a reproducible and scalable computational framework designed to decode the regulatory logic of 3D chromatin interactions through integrative analysis of HiChIP-derived enhancer-promoter interactions and transcription factor footprinting. By applying this framework to the human Middle Frontal Gyrus (MFG) and Substantia Nigra (SN), we demonstrate that functional specialization of the human brain is associated with region-specific enhancer-promoter wiring. Our results show that region-biased enhancer-promoter interactions are accompanied by coordinated differences in transcription factor programs, genetic risk enrichment, and evolutionary regulatory elements. In particular, MFG-biased interactions are enriched for neuropsychiatric risk variants and HARs, which are embedded within cortical regulatory interactions. Together with the elevated expression of associated neurodevelopmental genes in humans relative to non-human primates, these findings suggest that evolutionarily accelerated noncoding elements may have contributed to the emergence of human cognitive regulatory programs by reshaping cortical enhancer-promoter wiring.

The complexity of HiChIP data analysis, involving multi-step alignment, normalization, and statistical loop detection, has historically been hindered by fragmented workflows and implicit preprocessing assumptions. SnakeHichipTF addresses these challenges by formalizing the analytical logic of multiple engines within a unified Snakemake-based architecture. By locking dependencies within Conda environments and generating an automated provenance record, the framework ensures reproducible execution across computational systems, an increasingly critical requirement as 3D genomics expands into large-scale and translational studies. The inclusion of a Streamlit-based graphical interface further enhances accessibility, enabling researchers to interrogate spatial regulatory landscapes without extensive bioinformatics expertise while maintaining methodological rigor.

Our comparative analysis of the MFG and SN highlights functional partitioning of the 3D genome along distinct anatomical and physiological axes. A central contribution of this work is the integration of TF footprinting with chromatin interaction maps. While architectural proteins such as CTCF are broadly shared across regions, binding intensity and occupancy of additional TFs diverge between the MFG and SN. MFG-biased interactions preferentially recruit TFs associated with neuronal development and transcriptional activation, whereas SN-biased interactions are enriched for TFs linked to metabolic and stress-responsive regulation. These observations support a hierarchical model of chromatin organization in which common architectural scaffolds provide structural stability, while region-enriched TF programs modulate loop strength and enhancer engagement to generate functional specificity. In this framework, transcription factors act not only as downstream effectors but also as active organizers of region-specific 3D regulatory landscapes.

The integration of Human Accelerated Regions (HARs) into MFG-biased regulatory interactions further suggests an evolutionary dimension to cortical enhancer wiring. Our analyses indicate that HAR-overlapping anchors preferentially recruit MFG-enriched transcription factors and participate in region-biased enhancer-promoter interactions. Certain HAR-associated genes exhibit elevated expression in the human MFG relative to other primates. These findings are consistent with a model in which evolutionarily accelerated regulatory elements are incorporated into existing enhancer-promoter frameworks, contributing to human-specific transcriptional modulation in the prefrontal cortex.

A major challenge in the post-GWAS era is the “variant-to-function” bottleneck, linking noncoding risk variants to their target genes and regulatory mechanisms. By identifying enhancer-promoter interactions that harbor psychiatric risk loci and integrating TF binding inference, SnakeHichipTF provides a mechanistic bridge between genetic variation and spatial genome organization. For example, MFG-biased loops connecting distal enhancers to promoters of neuropsychiatric-associated genes offer a structural context for interpreting how noncoding variants may perturb regulatory networks. This approach moves beyond proximity-based gene assignment and instead situates risk variants within the three-dimensional nuclear environment.

Beyond the biological insights presented here, SnakeHichipTF establishes a reproducible computational infrastructure for linking chromatin architecture to transcription factor programs. By reconstructing the analytical logic of multiple HiChIP engines within a unified workflow and integrating footprint-based TF inference, the framework connects structural and mechanistic layers of gene regulation. Although this study focused on two brain regions, the modular design of SnakeHichipTF enables application to other tissues, developmental stages, and disease contexts. Future integration with single-cell chromatin profiling and perturbation-based validation will further clarify how transcription factor networks sculpt 3D genome organization across biological systems.

## Methods

### Pipeline architecture

SnakeHichipTF is implemented in Snakemake[35], providing a readable, scalable workflow that runs on local workstations, HPC clusters, and cloud platforms. All third-party software is distributed through Conda/Bioconda[36], eliminating manual dependency management and the need for administrator privileges. The pipeline was organized into discrete modules with explicitly defined inputs, outputs, logs, and benchmark files, thereby facilitating data provenance, reproducibility, resumable execution, and efficient parallelization.

### Genome setup

Reference genome FASTA files were parsed and filtered to retain only canonical chromosomes. For each supported organism, primary chromosomes (e.g., autosomes and sex chromosomes) were identified using organism-specific regular expression patterns, and noncanonical scaffolds and auxiliary contigs were excluded. The filtered genome was indexed using samtools faidx[37], and chromosome size files were derived from the resulting FASTA index. To support downstream interaction callers with distinct alignment backends, the workflow generated reference indices for both Bowtie2[38] and Chromap[17].

### Genomic feature generation and bias correction

A key requirement in HiChIP and PLAC-seq analysis is the accurate representation of restriction enzyme (RE) fragmentation and the associated sequence-derived biases that affect contact recovery. SnakeHichipTF includes an embedded configuration module supporting a broad range of restriction enzymes, including HindIII, MboI, NcoI, DpnII, BglII, MseI, HinfI, NlaIII, as well as multi-enzyme designs such as Arima. Because different interaction-calling frameworks rely on distinct statistical formulations for background correction, they also require different representations of genomic bias covariates, including fragment length, GC content, and sequence mappability. To standardize preprocessing across methods, SnakeHichipTF performs a unified in silico genome characterization step and then derives tool-specific feature files from the same underlying genome model.

Restriction fragments were first generated using the HiC-Pro[18] utility digest_genome.py, which produces a BED representation of adjacent RE-defined fragments. In parallel, genome-wide mappability was calculated using GenMap[39] with a default k-mer length of 50 and up to two mismatches, generating a high-resolution mappability track in BigWig format. These common baseline annotations were then reformatted according to the requirements of individual downstream tools.

For MAPS[7], which models background interactions in fixed-size genomic bins, the pipeline first quantified average mappability at RE fragment ends. Fragments with anomalous lengths (>20 kb) or low average mappability (<0.5) were excluded. The retained fragment-level features were then aggregated into user-specified genomic bins to generate the feature matrix required by the MAPS background model.

For FitHiChIP[19], which incorporates bias regression at the fragment or peak-pair level, SnakeHichipTF extracted 200-bp windows flanking both ends of each RE fragment and computed GC content using bedtools nuc and average mappability over the same windows. These values were combined into a BED-formatted feature track compatible with FitHiChIP.

For HiC-DC+[20], the workflow used the construct_features function implemented in the HiC-DC+ package to generate the native compressed bintolen covariate file. This step integrates effective fragment length, GC content, and mappability information directly from the in silico RE fragment model.

For hichipper[21], which primarily depends on accurate fragment boundary definitions, the workflow supplied the original RE-resolution fragment map generated during the unified digestion step.

By centralizing covariate generation and deriving all tool-specific inputs from the same genome model, SnakeHichipTF minimizes preprocessing inconsistencies and ensures that differences in interaction calls reflect algorithmic differences among statistical frameworks rather than differences in upstream feature preparation.

### HiChIP data processing and interaction quantification

SnakeHichipTF integrates four widely used HiChIP interaction-calling frameworks within a unified Snakemake architecture: MAPS[7], FitHiChIP [19], HiC-DC+[20], and hichipper[21]. Several of these tools were originally distributed as loosely connected shell scripts or script collections with limited fault tolerance and limited support for resumable execution. To improve robustness while preserving the original analytical logic of each method, we decomposed these workflows into environment-controlled Snakemake rules with explicit checkpoints and standardized parameter handling.

#### 1. Alignment and peak calling

Because different callers depend on different upstream alignment strategies, SnakeHichipTF provides two alignment branches.

For FitHiChIP [19], HiC-DC+[20], and hichipper[21], paired-end FASTQ files were processed using HiC-Pro[18] (v3.1.0). Reads were aligned with Bowtie2[38], valid ligation products were identified, and common Hi-C artifacts such as dangling ends, self-circles, and PCR duplicates were removed. Raw contact matrices and ICE-normalized matrices were then generated at the user-specified bin size. When requested, downsampling was applied at the valid-pair level, after which raw and ICE-normalized matrices were rebuilt from the subsampled valid pairs.

For MAPS[7], which is designed around a distinct preprocessing framework, reads were processed through a separate Chromap-based branch. Reads were aligned to the Chromap index[17], filtered using a user-defined mapping quality threshold, and deduplicated using an optical duplicate distance parameter.

To define active regulatory anchors for peak-centric interaction callers, one-dimensional enrichment peaks were identified using MACS2[40] from aligned HiChIP BAM files.

#### 2. Interaction calling with MAPS

For MAPS[7], Chromap-aligned reads were first processed using a feather_split preprocessing module from MAPS[7] to generate duplicate-filtered paired BAM files and long-range intra-chromosomal BEDPE contacts. These contacts were then combined with MACS2[40] peak calls and the precomputed bin-level genomic feature table. The resulting data were used in the MAPS[7] regression framework, which applies a regression-based background model to estimate expected interaction frequencies and assign statistical significance to observed contacts. Significant interactions were reported as BEDPE files at the selected bin resolution.

#### 3. Interaction calling with FitHiChIP

For FitHiChIP [19], the workflow was reconstructed as a series of explicit processing stages rather than a single monolithic execution step. Starting from HiC-Pro-derived contact matrices, the pipeline first generated initial interaction tables, then filtered cis interactions according to genomic distance thresholds. Coverage tracks were computed from peak-overlapping bins, and bias values were estimated using either coverage-based normalization or ICE-derived bias vectors, depending on the selected mode. These bias-corrected features were then joined to the interaction table, and interactions were sorted by genomic distance. Statistical significance was estimated separately for peak-to-all and all-to-all interaction modes, following the original FitHiChIP framework. Significant interactions were further merged across neighboring bins to recover clustered loop calls, and summary plots of contact count versus genomic distance were generated for both raw and merged significant interactions.

#### 4. Interaction calling with HiC-DC+

For HiC-DC+[20], HiC-Pro-derived valid pairs were analyzed together with the precomputed bintolen genomic covariate file. Significant interactions were identified using the native HiCDCPlus function from HiC-DC+, which models interaction counts under a negative binomial background model and reports significant contacts.

#### 5. Interaction calling with HiC-DC+ and hichipper

For hichipper[21], the workflow supplied HiC-Pro valid pairs, MACS2 peaks, and the RE fragment BED file to the standard hichipper loop-calling procedure. Distance-based filtering parameters were retained, including a minimum interaction distance of 5 kb and a user-defined upper distance threshold. Native hichipper outputs were standardized by renaming and relocating loop, anchor, and summary files into a consistent directory structure.

#### 6. Quality control and interoperability

SnakeHichipTF includes a dedicated QC module to summarize library complexity, alignment quality, and interaction characteristics across samples and callers. Metrics such as the fraction of reads in peaks (FRiP), valid-pair ratios, and interaction-distance quantiles were aggregated from the corresponding alignment and loop-calling modules and summarized using MultiQC[41]. To facilitate visualization and interoperability with downstream 3D genome tools, HiC-Pro matrices were automatically converted to.cool and.h5 formats using HiCExplorer[42]. Aligned BAM files were additionally converted to BigWig signal tracks where needed for downstream visualization and quantile-based quality assessment.

### Quantitative differential interaction analysis

To identify condition-specific enhancer interaction changes, SnakeHichipTF incorporates a differential interaction module based on hicdcdiff, which is distributed as part of the HiC-DC+[20] framework. Experimental design was specified through a sample sheet describing sample identity, condition labels, and replicate structure. The workflow automatically parsed this metadata to construct the comparison design and group replicate-level interaction files.

For statistical testing, interaction counts across candidate loops were modeled using a negative binomial distribution. Dispersion was estimated in a condition-aware manner using the user-specified fitType parameter, allowing the framework to accommodate the overdispersed count structure typical of sparse HiChIP data. The generalized linear modeling framework implicitly accounts for structural biases, including genomic distance and other systematic effects encoded in the HiC-DC+ model.

P values for differences in interaction strength between conditions were computed for all tested interactions and adjusted for multiple hypothesis testing using the Benjamini–Hochberg procedure. Interactions passing the specified false discovery rate threshold were retained as differentially interacting enhancer-promoter contacts for downstream integrative analyses.

### Transcription factor inference using scPrinter

To infer transcription factor occupancy within enhancer interaction regions, SnakeHichipTF integrates scPrinter[16], a deep learning–based framework for footprint-informed TF analysis from chromatin accessibility data. This module links HiChIP-derived regulatory structure with matched ATAC-seq signal to characterize TF programs associated with chromatin looping.

The workflow supports two input modes: aligned ATAC-seq BAM files and precomputed fragment files in compressed TSV format. For BAM-based input, reads were name-sorted using samtools, converted into paired-end fragment coordinates using bedtools[43] bamtobed-bedpe, and filtered to retain properly ordered intrachromosomal fragments. Fragment tables were then sorted, compressed with bgzip, and indexed with tabix.

scPrinter was run on the processed fragment files together with enhancer regions derived from HiChIP interaction anchors. These enhancer regions served as candidate regulatory elements for TF analysis. scPrinter was executed with GPU acceleration when available and generated TF binding scores across candidate regions together with summary tables and diagnostic scatter plots.

### Transcription factor inference using TOBIAS

In parallel, we applied TOBIAS[15] to matched ATAC-seq BAM files to perform motif-centered TF footprinting. Enhancer coordinates were derived from HiChIP interaction anchors and merged across samples to create a unified candidate region set. Depending on the input type, paired-end BAM files were either linked directly or converted from fragment-based representations.

First, Tn5 insertion bias was corrected using the ATACorrect module, which models both sequence bias and local accessibility. Corrected accessibility signals were generated as BigWig tracks. Next, FootprintScores was used to compute per-base footprint profiles across enhancer regions.

Motif instances were obtained either from user-supplied motif libraries or from default curated motif sets. Motifs were standardized and merged using TOBIAS Format Motifs. Differential TF binding activity across samples or conditions was then inferred using BINDetect, which integrates motif occurrence with corrected footprint signal. Bound sites identified for each TF were aggregated into BED files on a per-sample basis, and PlotAggregate was used to visualize average footprint profiles at predicted binding sites.

### Brain region-specific HiChIP datasets processing

We analyzed published H3K27ac HiChIP datasets from human Middle Frontal Gyrus (MFG) and Substantia Nigra (SN) generated within a single study (GEO: GSE147672). Because both brain regions were produced under the same experimental project and processing framework, direct comparison between MFG and SN minimizes confounding due to cross-study differences in sample handling, library preparation, and sequencing protocols.

All analyses were performed using our unified workflow SnakeHichipTF, which standardizes reference resources (BSgenome.Hsapiens.UCSC.hg38) and preprocessing steps across regions. For interaction calling and downstream differential analysis, we used HiC-DC+[20] outputs throughout. We focused on HiC-DC+ because it outputs statistically calibrated interaction calls and effect-size–like quantitative measures that are well suited for differential modeling, enabling direct comparison of interaction strength between MFG and SN under a unified framework. We processed both regions using identical parameters (genome build, bin size, filtering criteria, and significance thresholds), ensuring that observed regional differences reflect biological divergence rather than analytical inconsistencies.

### Gene expression analysis

Gene expression levels were quantified from brain regional RNA-seq data (GSE127898) and summarized as transcripts per million (TPM). Replicates and subregions belonging to the same brain region were combined prior to downstream analysis. Expression distributions of genes associated with enhancer interactions were compared between the two brain regions.

### GWAS variant enrichment analysis

To investigate the distribution of disease-associated genetic variation across regulatory architecture, we examined the overlap between enhancer interactions and genome-wide association study (GWAS) variants. GWAS data were obtained from the UCSC Genome Browser GWAS Catalog track[22] (mirroring the NHGRI-EBI GWAS Catalog; accessed November 2025). The dataset was processed into BED format, with columns specifying genomic coordinates (chromosome, start, end) and the reported trait in the fourth column. This processing yielded 713,013 unique single-nucleotide polymorphisms (SNPs) linked to 31,944 distinct reported traits (as originally described in the source publications, prior to EFO ontology standardization). SNPs were intersected with enhancer interactions across brain regions. Enrichment analyses were performed at the enhancer-bin level using fisher test, stratified by hub class. Only associations with a p-value ≤ 0.01 and an odds ratio ≥ 5 were retained for further analysis.

### Gene Ontology enrichment analysis

GO enrichment analyses were performed using Enrichr (v3.4)[44]. Analyses were conducted separately for region-biased TFs.

## Data visualization

The box plots, barplots, donut plots and radar chart were produced using ggplot2 (v.3.5.2; https://ggplot2.tidyverse.org), heat maps were produced using pheatmap (v.1.0.13; https://cran.r-project.org/web/packages/pheatmap/index.html) in R (v.4.4.3). The representative tracks were produced using PygenomeTracks (v3.9)[45]. Aggregation plots were generated using hicAggregateContacts from HiCExplorer[42] (v3.7.6).

## Data availability

H3K27ac HiChIP and ATAC-seq data from human brain regions were obtained from GEO under accession number GSE147672. Brain region RNA-seq data were also obtained from GEO under accession number GSE127898.

## Code availability

SnakeHichipTF snakemake pipeline is publicly available on GitHub at https://github.com/YidanSunResearchLab/SnakeHichipTF.git

## Acknowledgements

We thank all members of our department and institutes for fostering a collaborative and supportive research environment.

## Authors’ contributions

J.T. and Y.S. conceived the study. J.T. and Y.W. developed the computational framework, performed data analyses, and generated all figures. R.H. contributed to the pipeline design and interpretation of the brain region–specific analyses. Y.S. supervised the project, guided methodological development, and contributed to data interpretation. J.T. and Y.S. drafted the initial version of the manuscript, and J.T., Y.W., R.H. and Y.S. jointly revised and finalized the manuscript.

## Ethics declarations Competing interests

The authors have declared that no competing interests exist.

## Supplementary information

Supplementary Table 1: Differential enhancer interactions analysis between two brain regions.

